# Transcriptomic signature and metabolic programming of bovine classical and nonclassical monocytes indicate distinct functional specializations

**DOI:** 10.1101/2020.10.30.362731

**Authors:** Stephanie C. Talker, G. Tuba Barut, Reto Rufener, Lilly von Münchow, Artur Summerfield

## Abstract

Similar to human monocytes, bovine monocytes can be split into CD14^+^CD16^−^ classical and CD14^−^CD16^+^ nonclassical monocytes (cM and ncM, respectively). Here, we present an in-depth analysis of their steady-state transcriptomes, highlighting pronounced functional specializations. Gene transcription indicates that pro-inflammatory and antibacterial processes are associated with cM, while ncM appear to be specialized in regulatory/anti-inflammatory functions and tissue repair, as well as antiviral responses and T-cell immunomodulation. In support of these functional differences, we found that oxidative phosphorylation prevails in ncM, whereas cM are clearly biased towards aerobic glycolysis. Furthermore, bovine monocyte subsets differed in their responsiveness to TLR ligands, supporting an antiviral role of ncM. Taken together, these data clearly indicate a variety of subset-specific functions in cM and ncM that are likely to be transferable to monocyte subsets of other species, including humans.

## 1 Introduction

With their high functional plasticity (1, 2), monocytes are a central component of the mononuclear phagocyte system (MPS). Although their delineation from *bona fide* macrophages and *bona fide* dendritic cells has proofed challenging, especially in tissues, monocytes and monocyte-derived cells are now fully appreciated as a separate lineage (2–4). Monocytes in blood of humans and cattle can be subdivided into at least two different subsets based on the expression of CD14 and CD16 (5–7): classical monocytes (cM; CD14^+^CD16^−^) and nonclassical monocytes (ncM; CD14^−^CD16^+^). In mice, analogous subsets can be defined by Ly6C expression (8). A third intermediate monocyte subset (intM; CD14^+^CD16^+^) is less well defined and has been shown to transcriptionally resemble ncM in both cattle and human (5, 6, 9). It is widely accepted that murine and human intM and ncM arise by sequential differentiation from cM in the periphery (10–12), with development and survival of ncM depending on the transcription factor NR4A1 (13). In both human and cattle, cM are described as the dominant subset, comprising about 80% of all monocytes in peripheral blood, while ncM and intM comprise only small fractions (about 10% each) (2, 7). In mice, however, ncM (Ly6C^−^) are reported to be as frequent as cM (Ly6C^+^) in peripheral blood (8).

Monocytes in general have been described to be capable of antigen presentation to T cells, however whether they are as potent as dendritic cells remains controversial (1). Notably, TLR7 stimulation, but not TLR3 or TLR4 stimulation, has been shown to promote cross-presenting abilities in murine Ly6C^+^ monocytes (14). Also bovine monocytes, particularly ncM, were reported to induce allogeneic T-cell responses (15). However, neither resting nor TLR-stimulated bovine monocytes express CCR7 (16), which is required for migration of antigen presenting cells from the site of inflammation to the draining lymph node. Classical monocytes are known for their pro-inflammatory role especially in bacterial infections (17), however the role of intM and ncM is less well described. Nonclassical monocytes are generally viewed as anti-inflammatory and vasoprotective (18). In fact, ncM have been termed “housekeepers of the vasculature”, as they were found to crawl along vascular endothelium (8, 19) and sustain vascular integrity by orchestrating endothelial renewal (20). In response to TLR7 stimulation, murine ncM have been shown to recruit neutrophils to the endothelium and to clear neutrophil-induced focal necrosis (20). This may indicate that ncM play a pivotal role in antiviral immunity.

Also in humans, ncM were shown to patrol blood vessels and to be specialized in sensing nucleic acids via TLR7 and TLR8 (21). The transcription of molecules for endothelial adhesion in bovine ncM (7, 9) supports a similar role in cattle.

Furthermore, monocyte-derived macrophages have been implicated in wound healing and tissue regeneration (22, 23), with murine ncM described as biased progenitors of wound healing macrophages (24). This is in line with an earlier study, suggesting sequential recruitment of cM and ncM, rather than *in-situ* conversion of cM into ncM, during the inflammatory and healing phase of myocardial infarction (25).

Recent transcriptomic analyses of bovine monocyte and dendritic-cell subsets performed by our group have confirmed the pronounced differences between cM and ncM, and suggest that bovine intM are transcriptionally almost identical to ncM (9). We have reanalyzed the transcriptomic dataset in greater detail, this time focusing on monocytes only, and have performed intensive literature research on differentially expressed genes in order to get insights into the functional specializations of bovine monocyte subsets. These in-depth analyses have revealed subset-specific gene expression indicating opposing and complementary functions in acute inflammation, antibacterial and antiviral responses, as well as T-cell responses, resolution of inflammation and tissue repair. Our analyses of metabolism and TLR responsiveness support the idea that bovine cM and ncM are functionally highly specialized subsets that deserve to be studied in greater detail in the future, in the context of infection and vaccination.

## 2 Materials and Methods

### 2.1 Isolation of bovine PBMC

Blood of cows was collected at the Clinic for Ruminants (Vetsuisse Faculty, University of Bern, Bern, Switzerland) by puncturing the jugular vein, using citrate-based Alsever’s solution (1.55 mM of C_6_H_12_O_6_, 408 mM of Na_3_C_6_H_5_O_7_∙2H_2_O, 1.078 mM of NaCl, and 43 mM of C_6_H_8_O_7_, pH 6.2) as an anticoagulant. Blood sampling was performed in compliance with the Swiss animal protection law and approved by the animal welfare committee of the Canton of Bern, Switzerland (license numbers BE102/15 and BE104/17). For peripheral blood mononuclear cell (PBMC) isolation, blood was centrifuged at 1000 × *g* for 20 min (20 °C), the buffy coat was collected, diluted with PBS to a ratio of 1 to 1 (room temperature), and layered onto lymphocyte separation medium (1.077 g/ml; GE Healthcare Europe GmbH, Freiburg, Germany). After centrifugation (800 × *g*, 25 min, 20°C), PBMC were collected and washed twice (400 × *g*, 8 min, 4°C) with cold PBS containing 1 mM UltraPure^TM^ EDTA (Invitrogen, ThermoFisher, Basel Switzerland). In order to remove platelets, a final washing step was done at 250 × *g* (8 min, 4°C).

### 2.2 Phenotyping of monocyte subsets by flow cytometry

Phenotyping of bovine monocyte subsets was performed with freshly isolated PBMC in 96-well U-bottom microtiter plates (1 × 10^7^ cells per sample). Antibodies used for the two-step five-color stainings are shown in Table 1. Incubations were performed for 20 min at 4 °C. Washing steps between incubations (400 × *g*, 4 min, 4 °C) were done with Cell Wash (BD Biosciences). Prior to staining, PBMC were incubated with bovine IgG in order to block Fc receptors (ChromPure mouse IgG, Jackson ImmunoResearch). For detection of dead cells, Live/Dead Near-IR stain (ThermoFisher) was included in the last incubation step. Compensation was calculated by FACSDiva software using single-stained samples. For each marker to be examined on monocyte subsets, a fluorescence-minus-one (FMO) control was included. Samples were acquired with a FACSCanto II flow cytometer (BD Biosciences) equipped with three lasers (405, 488, and 633 nm). At least 5 × 10^5^ cells were recorded in the “large-cell” gate.

**Table 1:** Antibodies used for flow-cytometric phenotyping of monocyte subsets.

| Panel 1 |  |  |  |
| --- | --- | --- | --- |
|  | Antigen | Clone / Source of mAb | Detection /Source |
| Core | CD14 | CAM66A / Kingfisher | Anti-IgM-PE / Southern Biotech |
|  | CD16 | KD1 / Bio Rad | Anti-IgG2a-AF647 / Molecular Probes |
|  | CD172a | CC149 / Bio Rad | Anti-IgG2b-AF488 / Molecular Probes |
| Phenotypic marker (Pm) | CD5 | AFRCIAH-CC29 / in house | Anti-IgG1- BV421 / BioLegend |
|  | CD8α | CACT80C / Kingfisher |  |
|  | CD11a | HUH73A / Kingfisher |  |
|  | CD40 | IL-A156 / Kingfisher |  |
|  | CD43 | CO.44B8 / Bio-Rad |  |
|  | CD45 | 1.11.32 / Bio-Rad |  |
|  | CD62L | Du-1-29 / in house |  |
|  | CD80 | IL-A159 / Kingfisher |  |
|  | CD86 | IL-A190A / Kingfisher |  |
|  | CD163 | LND68A / Kingfisher |  |
|  | CD205 | IL-A114 / Bio Rad |  |
|  | MHC-II | VPM54 / in house |  |
| Panel 2 |  |  |  |
|  | Antigen | Clone / Source of mAb | Detection /Source |
| Core | CD14 | CAM36A / Kingfisher | Anti-IgG1-BV421 / BioLegend |
|  | CD16 | KD1 / Bio Rad | Anti-IgG2a-AF647 / Molecular Probes |
|  | CD172a | CC149 / Bio Rad | Anti-IgG2b-AF488 / Molecular Probes |
| Pm | CD11c | BAQ153A / Kingfisher | Anti-IgM-PE / Southern Biotech |
| Panel 3 |  |  |  |
|  | Antigen | Clone / Source of mAb | Detection /Source |
| Core | CD14 | CAM66A / Kingfisher | Anti-IgM-PE / Southern Biotech |
|  | CD16 | KD1 / Bio Rad | Anti-IgG2a-AF647 / Molecular Probes |
|  | CD172a | DH59B / Kingfisher | Anti-IgG1-BV421 / BioLegend |
| Pm | CD11b | MM10A / Kingfisher | Anti-IgG2b-AF488 / Molecular Probes |

### 2.3 Fluorescence-activated cell sorting and RNA sequencing

Illumina sequencing data shown in the present manuscript are derived from previous experiments, described in Talker et al. (9). Experimental procedures are therefore described only briefly and the reader is referred to our previous publication for more details (9). In order to sort bovine monocyte subsets for Illumina sequencing, a two-step staining was performed with 3 × 10^8^ freshly isolated PBMC as reported previously (9). Classical monocytes were sorted as CD172a^high^CD14^high^CD16^−^, intM as CD172a^high^CD14^high^CD16^high^ and ncM as CD172a^high^CD14^−^CD16^high^ using a FACS Aria (BD Biosciences). All sorted subsets had a purity of at least 97%. Per subset, at least 100,000 sorted cells were frozen in TRIzol (ThermoFisher) for later RNA extraction (Nucleospin RNA kit, Macherey Nagel). High-quality RNA (approximately 500 ng; RIN 8) was used for nondirectional paired-end mRNA library preparation (TruSeq Sample Preparation Kit; Illumina).

Sequencing was performed on the Illumina HiSeq3000 platform using 100 bp single-end sequencing, yielding between 25.2 and 41.1 million read pairs per sample. Reads were mapped to the *Bos taurus* reference genome (UMD3.1) with Hisat2 v. 2.1.0, and FeatureCounts from Subread v. 1.5.3 was used to count the number of reads overlapping with each gene, as specified in the Ensembl annotation (release 91). The RNA-seq data are available in the European Nucleotide Archive (http://www.ebi.ac.uk/ena) under the accession number PRJEB28324. Raw counts of the sequencing data previously published in Talker et al. 2018 (9) were re-analyzed with the Bioconductor package DESeq2 (26), including only data for the three monocyte subsets and considering the factor “animal” in the design formula. Raw counts were normalized to account for differences in sequencing depth between samples. Gene length was not considered. Lists of differentially expressed genes were obtained by performing pairwise comparisons with DESeq2 and by filtering genes with a p-adjusted value of 0.05. Gene lists were manually screened for genes of interest using the human gene database GeneCards^®^ and literature research via PubMed^®^.

Principal component analysis was performed with normalized and rlog-transformed counts of the 1000 most variable genes across monocyte samples. Heatmaps were prepared following log2-transformation of normalized counts. Prior to log2 transformation, a pseudocount of 1 was added to the values to avoid zeros. All analyses were performed using R version 3.6.1.

### 2.4 Extracellular flux analysis

For metabolic Seahorse assays, bovine monocyte subsets were FACS-sorted (Moflo Astrios EQ) from magnetically pre-sorted CD172a^+^ PBMC. Combined flow cytometry and magnetic staining was performed in 50 mL falcon tubes and included five incubation steps, each carried out at 4°C for 20 min. Staining, and washing (400 × g, 8 min, 4°C) in-between each incubation step was performed in PBS containing 1 mM EDTA and 5 % (v/v) heat-inactivated FBS (GIBCO, Life Technologies, Basel, Switzerland). In a first step, freshly isolated PBMC (4 × 10^8^) were incubated with bovine IgG to block Fc receptors. This was followed by incubation with the primary antibodies anti-CD172a (CC149, IgG2b), anti-CD14 (CAM36A, IgG1), and anti-CD16 (KD1, IgG2a) and the secondary antibodies anti-IgG1-AF647 (Molecular Probes) and anti-IgG2a-PE (Southern Biotech). In a forth step, anti-mouse IgG magnetic beads (Miltenyi Biotec) were added. Following this, cells were loaded onto two LS columns (Miltenyi Biotec) for magnetic enrichment of CD172a expressing cells. In a fifth step, anti-IgG2b-AF488 (Molecular Probes) was added, resulting in a dim staining of CD172a for FACS. Enriched monocytes were sorted on a MoFlo Astrios EQ cell sorter equipped with five lasers (Beckman Coulter Eurocenter SA, Nyon, Switzerland) at the flow cytometry core facility of the University of Bern. Purity of subsets was confirmed by re-analysis of samples and was shown to be at least 98%. Following sorting, cells were resuspended in pH-optimized Seahorse assay medium (Seahorse XF DMEM Medium supplemented with 10 mM Seahorse XF Glucose, 1mM Seahorse XF Pyruvate, and 2mM Seahorse XF Glutamine) and 5 × 10^5^ cells (*ex vivo*) or 1 × 10^5^ cells (after 6-day culture) were seeded in duplicates or triplicates into an 8-well Seahorse plate (Seahorse XFp FluxPak). Two wells served as background controls. During manual cell counting, viability of sorted subsets was confirmed to be 100% by trypan blue staining. Cells not immediately used for Seahorse assays, were incubated for six days at 37°C to allow for differentiation into monocyte-derived macrophages. Specifically, remaining cells were seeded into a 12-well plate with 1 × 10^6^ cells per well in 2 ml culture medium consisting of DMEM-GlutaMAX^TM^ (Gibco, ref.no: 31966-021), supplemented with penicillin (50 I.U./mL), streptomycin (50 μg/mL), 10% heat-inactivated FBS, and 1% M-CSF (produced in house; titrated to optimize viability of monocytes in culture). After six days of incubation, cells were harvested, resuspended in Seahorse assay medium, counted, and processed as described above. Viability (trypan blue) of harvested cells was above 90 %.

Seahorse preparations and measurements were performed according to the manufacturer’s instructions. For measurements with the Agilent Seahorse XFp Analyzer, the standard settings of the XFp Real-Time ATP Rate Assay were used. Following basal measurements, oligomycin (Sigma-Aldrich, cat. no. O4876; final concentration 9 μM), an inhibitor of ATP synthase (complex V of the ETC) was added. After another measurement phase, a mastermix of rotenone (Sigma-Aldrich, cat. no. R8875; final concentration 8 μM), an inhibitor of complex I, and antimycin A (Sigma-Aldrich, cat. no. A8674; final concentration 8 μM), an inhibitor of complex III, was added. Inhibitors of the electron transport chain were purchased from Sigma-Aldrich (St. Gallen, Switzerland) and titrated on bovine cM to determine optimal concentrations. For establishment of the procedure, Agilent Seahorse XFp Real-Time ATP Rate Assay Kit (cat. no. 103591-100) was used according to manufacturer’s instructions, but with an increased final concentration (6 μM) of the provided oligomycin, according to prior titration on bovine cM. Data were analyzed using the Seahorse Wave Desktop Software version 2.6.1 (Agilent).

### 2.5 Phosphoflow cytometry

The following TLR ligands were used at a final concentration of 10 μg/mL to assess TLR responsiveness of bovine monocyte subsets: Pam2CSK4 [C65H126N10O12S,3TFA] (InvivoGen, Labforce, Basel, Switzerland), high-molecular-weight (HMW) polyinosinic-polycytidylic acid [Poly(I:C)] (InvivoGen), LPS from E. coli strain K12 [LPS-EK] (InvivoGen), Gardiquimod (Sigma-Aldrich), Resiquimod (Sigma-Aldrich).

Stimulation with TLR ligands and staining for phosphorylated p38 MAPK was performed as reported previously (16). Prior to stimulation, cell surface staining was performed. Following Fc-receptor blocking with bovine IgG, defrosted and CD3-depleted bovine PBMC were stained with anti-CD172a (CC149, IgG2b), anti-CD14 (CAM36A, IgG1), and anti-CD16 (KD1, IgG2a), followed by incubation with anti-IgG2b-AF488 (Molecular Probes), anti-IgG1-biotin (Southern Biotech) and anti-IgG2a-PE. In a fourth step, mouse IgG (Jackson ImmunoResearch) was added together with Streptavidin-BV421 (BD Biosciences) and Live/Dead™ Fixable Near-IR stain (Thermo Fisher Scientific, Basel Switzerland). Stained cells were incubated for 15 minutes with respective TLR ligands in PBS or with PBS alone (waterbath at 37°C). Immediately after this incubation period, cells were fixed with BD Cytofix/Cytoperm™ fixation buffer (BD Biosciences; 12 min at 37 °C) and thereafter stained with anti-p38 MAPK-AF647 (BD Phosflow; 30 min at 37 °C). Samples were acquired with a FACSCanto II flow cytometer (BD Biosciences) equipped with three lasers (405, 488, and 633 nm). At least 1.5 × 10^6^ cells were recorded in the “large-cell” gate. Compensation was calculated by FACSDiva software using single-stained samples.

### 2.6 Preparation of figures

Figures were prepared using FlowJo version 10 (FlowJo LLC, Ashland, OR), GraphPad Prism version 7.03 for Windows (GraphPad Software, San Diego, CA), R version 3.6.1., and Inkscape (www.inkscape.org).

## 3 Results

### 3.1 Phenotype and transcriptome of nonclassical and intermediate monocytes differ markedly from classical monocytes

Bovine monocyte subsets were analyzed by flow cytometry for expression of various surface molecules (Figure 1A+B). While some molecules showed clear subset-dependent expression patterns (e.g. CD163, CD11a, CD11b, CD205), other molecules varied quite strongly with the animals analyzed (e.g. CD40, CD80, MHC-II). A gradual increase from cM over intM to ncM could be observed for expression of CD11a, whereas a gradual decrease was observed for CD11b. In addition to CD11a, ncM expressed the highest levels of CD5, CD8α, and CD205. Intermediate monocytes expressed the highest levels of CD86 and, for 2 out of 3 animals, also expressed the highest levels of CD40 and MHC-II. Highest expression on intM combined with lowest expression on ncM was seen for CD163 and CD11c. Expression of CD172a, CD45 and CD43 was increased in ncM and intM compared to cM. Furthermore, cM expressed the highest levels of CD62L, being almost absent from intM and ncM. Taken together, our phenotypic analyses have confirmed previously described expression patterns of MHC-II, CD172a, CD62L, CD11a and CD11b on bovine monocyte subsets (7) and provide new information on the expression of CD45, CD43, CD5, CD40, CD80, CD86, CD11c, CD163, and CD205. Moreover, in contrast to a previous study, where CD8α expression was concluded to be absent from all monocytes (27), we found indications of weak CD8α expression on ncM and intM.

**Figure 1.**
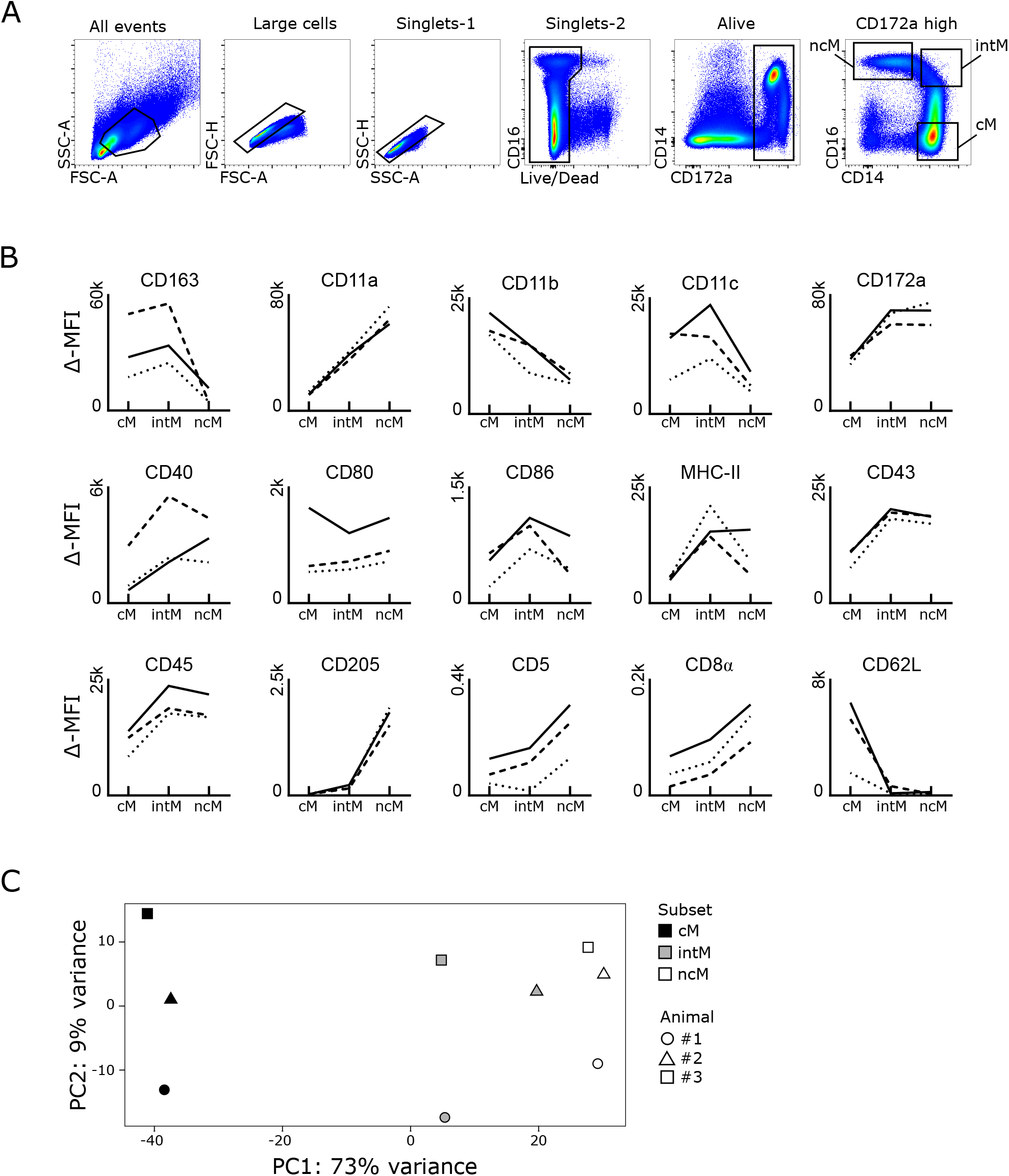
Phenotype and transcriptional clustering of bovine monocyte subsets. (A+B) Flow cytometric analysis. Freshly isolated PBMC were stained for flow cytometry. Monocyte subsets were gated based on expression of CD 14 and CD 16 within CD172a^high^ cells following gating on large cells (FSC^high^), single cells (within diagonal in FSC-A vs. FSC-H and SSC-A vs. SSC-H), and living cells (Near-IR^low^). Classical monocytes (cM) were gated as CD14^high^CD16^−^, intermediate monocytes (intM) as CD14^h,gh^CD16^high^ and nonclassical monocytes (ncM) as CD14^high^CD16^high^. (B) Graphs show the delta median fluorescence intensity (MFI) of surface expression for selected molecules. Delta MFI was calculated as the difference in MFI between stained samples and FMO controls. Lines illustrate the delta MFI across monocyte subsets. In total, data of seven different animals is shown. Within single graphs, data of three different animals is illustrated by solid, dashed, and dotted lines. (C) First two axes of a principle component analysis (PCA) including the 1000 most variable genes. Illumina sequencing was performed on RNA isolated from sorted monocyte subsets of three animals. Each dot represents one sample, with the color coding for different cell subsets and the shape coding for the three different animals.

Principal component analysis of previously published RNA-sequencing data (9) revealed that differences between monocyte subsets explained the highest proportion of total variance (73%, PC1) in the transcriptomic dataset, followed by 9% of variance (PC2) explained by differences between animals (Figure 1C). As expected from previous analyses (9), ncM and intM clearly clustered apart from cM, with intM clustering much closer to ncM than to cM. Notably, when looking at PC1, intM of animal #2 clustered closer to ncM samples than to intM samples of the other two animals.

As illustrated in Supplementary Figure 1A, the identity of monocyte subsets was confirmed by the subset-specific transcription of the key genes *NR4A1* (ncM, intM), *CXCR3* (ncM, intM) and *CCR2* (cM). Moreover, the transcription of surface molecules previously analyzed by flow cytometry in different animals followed the same patterns. Except for *CD163* and *ITGAM* (CD11b), for which mRNA content in intM was lower than expected (Supplementary Figure 1B). Pairwise comparisons of monocyte subsets revealed a variety of differentially expressed genes, involved in various different immune functions. The genes addressed in the following chapters have been selected based on pairwise comparisons and literature research and raise no claim for completeness.

### 3.2 Pro-inflammatory gene expression prevails in classical monocytes

Monocyte subsets clearly differed in the transcription of inflammatory cytokines and cytokine receptors (Figure 2A). Classical monocytes were strongly enriched in transcripts for IL-1 (*IL1A*, *IL1B*) and for the IL1-receptor (*IL1R1*, *IL1RAP*). Moreover, cM predominantely expressed *IL6R*, *IL15RA*, *IL17RA*, *IL17RC*, *IL17RD* and *IL27RA*. Trans-signaling with soluble IL-6 receptor is reported to mediate pro-inflammatory functions of IL-6 (28) and also *in vitro* stimulation with IL-27 is reported to increase inflammasome activation in monocytes (29). Furthermore, expression of *TNFSF13* (APRIL), reported to induce IL-8 production in the human macrophage-like cell line THP-1 (30) and *TNFRSF1A* (TNFR1), mediating pro-inflammatory signaling of TNF-α, was highest in cM. Notably, *TNF* was found to be primarily expressed in ncM and intM. Tumor necrosis factor alpha (*TNF*) is regarded as a master regulator of inflammation with various effects on different cell types. TNF-α is produced either in soluble form which signals mainly via TNFR1 (*TNFRSF1A*), or as transmembrane protein, being the main ligand for TNFR2 (*TNFRSF1B*) (31) and implicated in reverse signaling (32), the functional consequences of which are incompletely understood (32, 33). Moreover, receptors for IL-12 and IL-20 (*IL12RB1*, *IL12RB2* and *IL20RA*, *IL20RB*), were mainly expressed in ncM, though reads for *IL20RB* were very low (mean = 20). Although little is known about IL-12-receptor and IL-20-receptor signaling in monocytes, *in vitro* studies suggest overall pro-inflammatory effects (34, 35).

**Figure 2.**
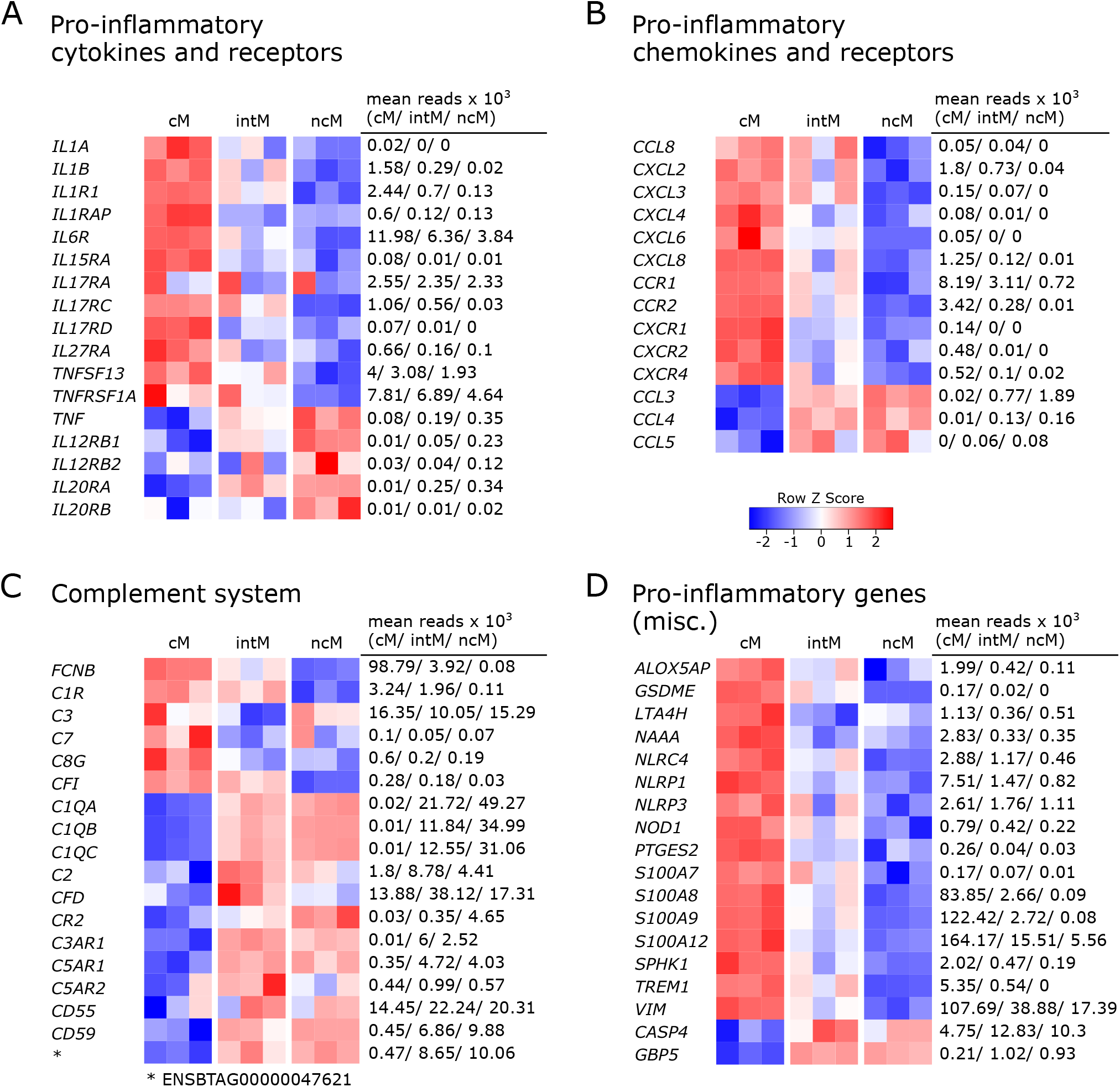
Pro-inflammatory gene expression. Illumina sequencing was performed on RNA isolated from sorted monocyte subsets (cM, intM, ncM) of three animals. Heatmaps show row z-scores calculated from log2-transformed normalized counts of selected genes coding for pro-inflammatory cytokines and receptors (A), pro-inflammatory chemokines and receptors (B), proteins associated with the complement system (C), and other pro-inflammatory mediators (D). Mean kilo reads for each subset and gene are given to the right of each heatmap.

Also a number of pro-inflammatory chemokines and chemokine receptors were found to be differentially expressed among monocyte subsets (Figure 2B). Looking at chemokines overexpressed in cM, the most pronounced differences between cM and ncM were found in the expression of *CXCL2* and *CXCL8*. Moreover, transcripts for *CCL8*, *CXCL3*, *CXCL4* and *CXCL6* could be detected in cM and in intM of two animals, though all with low number of reads.

Chemokine receptors associated with inflammation were all predominantly expressed by cM. Expression of *CCR1* in cM was upregulated 2-fold over intM and 11-fold over ncM. Expression of *CCR2* and *CXCR4* in cM was clearly upregulated over intM (12-fold and 5-fold, respectively), and almost absent from ncM. Exclusive expression in cM was observed for *CXCR1* and *CXCR2*. Nonclassical monocytes and intM clearly showed the highest expression of *CCL3* and were also enriched in transcripts for *CCL4* and *CCL5*, though at a lower level.

Gene expression also supports a prominent role of cM in complement-mediated inflammatory processes (Figure 2C). We found that transcription of *FCNB*, a recognition receptor for the lectin complement pathway, was highly increased in cM and almost absent in ncM, as was the transcription of *C1R*, a subunit of the complement C1. Notably, transcription of complement component *C3*, which is central for activation of both the classical and the alternative complement pathway, was enriched in cM and ncM, and showed the lowest transcription in intM, whereas complement factor I (*CFI*, C3b-Inactivator), an important negative regulator of both complement pathways, was exclusively transcribed in cM and intM. Moreover, transcription of *C7* as well as of *C8G*, both involved in formation of the membrane attack complex, was highest in cM. Nevertheless, certain important genes of the complement system were overexpressed in both ncM and intM. Among those genes were *C1QA*, *C1QB*, and *C1QC*, all of which were barely expressed in cM. The molecule C1q is one of the main sensors of PAMPs and DAMPs, but also antibody complexes, in the classical complement pathway and has been associated with tolerogenic functions (36). Notably, C1q also binds to the surface of dead cells, thereby promoting their phagocytosis. Complement factor 2 (*C2*) and complement factor D (*CFD*), involved in the classical and alternative complement pathway, respectively, showed the highest transcription in intM. The receptor for complement factor C3a (*C3AR1*) was exclusively expressed in intM and ncM, whereas transcription of the receptor for complement factor C3d (*CR2*/CD21) was markedly increased in ncM and to a lesser extent in intM, when compared to cM. Both known receptor genes for complement factor C5a, *C5AR1* (CD88) and *C5AR2* were increased in ncM and intM, with the latter showing the highest expression in intM. While C5aR1 is regarded as a mediator of pro-inflammatory signaling, C5aR2 has recently received attention as a multifaceted modulator of C5a signaling, described to dampen inflammasome activation and to alter TLR signaling (37). Furthermore, we found a higher transcription of *CD55* in ncM/intM compared to cM, a complement inhibitory protein reported to suppress T-cell responses (38). A 20-fold increased transcription in ncM as compared to cM was found for *CD59*, a receptor for C8 and C9. Apart from its function as an inhibitor of the membrane attack complex, CD59 expression on antigen-presenting cells was reported to deliver suppressive signals to murine CD59-expressing CD4 T cells via a complement-independent ligand (39, 40). Nonclassical monocytes were also clearly enriched in transcripts for the gene ENSBTAG00000047621. A protein query revealed the highest similarity with human (53.7%) and murine (47.5%) C4BPA, a molecule implicated in the inhibition of classical complement activation that has been reported to induce an anti-inflammatory state in monocyte-derived dendritic cells (41). However, no transcripts were found for the gene annotated as bovine C4BPA.

Also transcription of other genes associated with pro-inflammatory functions was clearly dominant in cM (Figure 2D). This includes genes coding for sensory components of inflammasomes (*NOD1*, *NLRC4*, *NLRP1*, *NLRP3*), pyroptosis-mediating gasdermin E (*GSDME*), enzymes involved in the biosynthesis of leukotriens (*ALOX5AP*, *LTA4H*) and prostaglandins (*PTGES2*), as well as the kinase for generation of sphingosine-1-phosphate (*SPHK1*), the pro-inflammatory receptor TREM1, the pro-inflammatory amidase NAAA (42), and S100 proteins promoting inflammation (*S100A7*, *S100A8*, *S100A9*, *S100A12*) (43). Furthermore, transcripts for vimentin (*VIM*), reported to be a key positive regulator of the NLRP inflammasome (44), were clearly enriched in cM. While none of the monocyte subsets expressed nitric-oxide synthases at steady state (data not shown), we could recently show that bovine cM massively increase transcription of *NOS2* upon *in vitro* stimulation with TLR ligands (16). Nonclassical monocytes expressed the highest levels of *CASP4* and *GBP5*, the latter being described as an activator of inflammasome assembly (45). Caspase 4 (*CASP4*), being part of the non-canonical inflammasome, is described to promote pro-inflammatory cytokine production, but recently has also been implicated in autophagy (46) – a process that may negatively regulate inflammasome signaling.

Taken together, the differential expression of pro-inflammatory genes suggests fundamentally different functions of bovine monocyte subsets.

### 3.3 Gene expression and TLR responsiveness indicate complementary functions of cM and ncM in antibacterial and antiviral immunity

Looking at the gene expression of pattern recognition receptors, we found that *TLR2* was expressed higher in cM and intM compared to ncM, and that *TLR4* and *TLR5* transcripts were clearly enriched in cM (Figure 3A). Toll-like receptor 6 (*TLR6*) showed a trend towards higher expression in intM and ncM. Expression of *TLR3* differed markedly between animals, but was in tendency highest in intM. Furthermore, cM expressed the highest levels of *TLR7* and *STING1*, the latter encoding a cytoplasmic receptor for DNA of both viral and bacterial origin. For two out of 3 animals, *TLR9* expression was also highest in cM. Transcript levels for RIG-1 (*DDX58*) and MDA-5 (*IFIH1*), however, were highest in ncM and intM.

**Figure 3.**
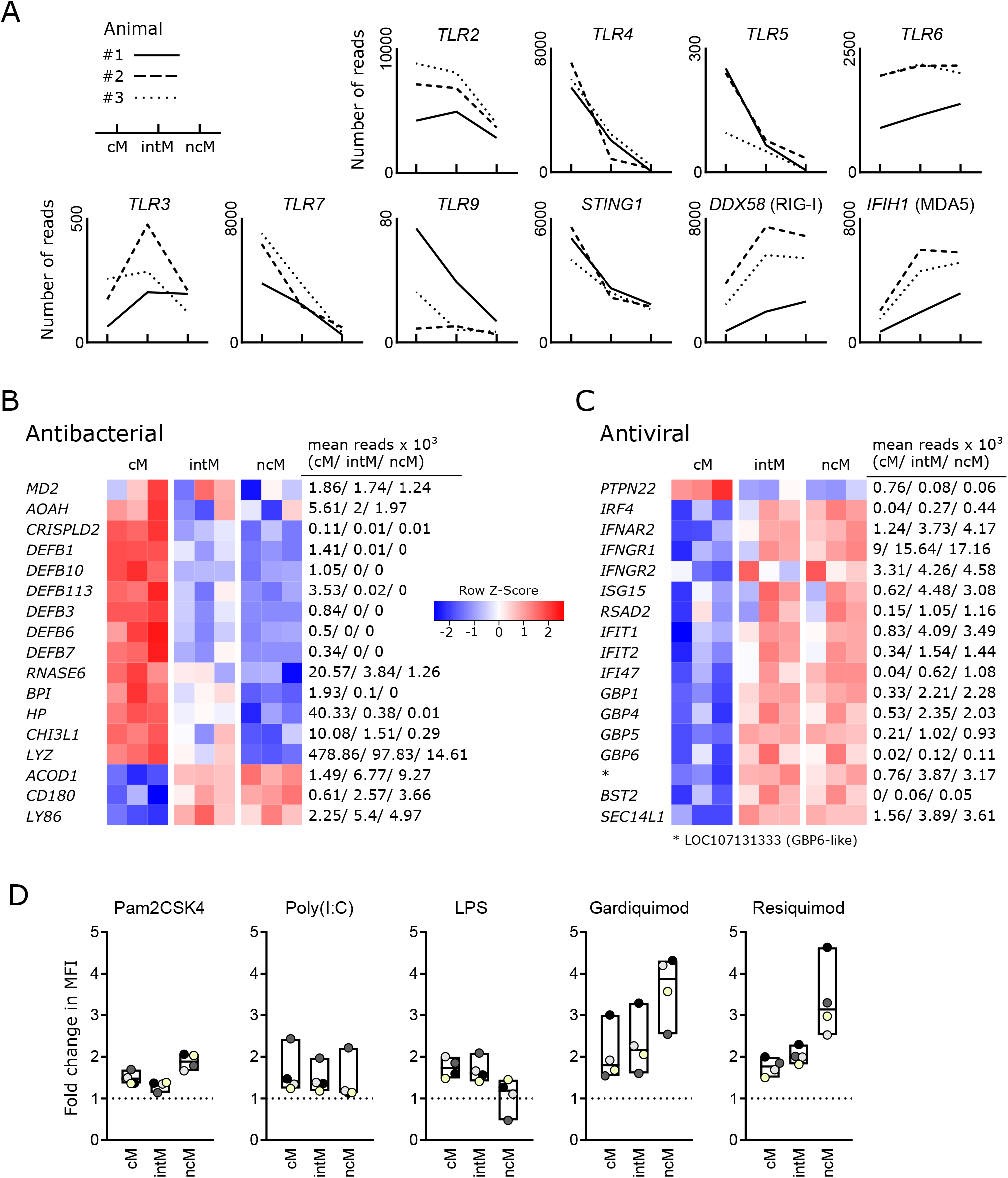
Antimicrobial gene expression and TLR responsiveness. Illumina sequencing was performed on RNA isolated from sorted monocyte subsets (cM, intM, ncM) of three animals. (A) Gene expression for pathogen-recognition receptors. Graphs show the number of reads across monocyte subsets for selected genes with individual animals indicated by solid (#1), dashed (#2) and dotted (#3) lines. (B+C) Heatmaps show row z-scores calculated from log2-transformed normalized counts for genes associated with antibacterial (B) and antiviral (C) responses. Mean kilo reads for each subset and gene are given to the right of each heatmap. (D) Responsiveness of bovine monocyte subsets to TLR ligands. Defrosted bovine PBMC were depleted of CD3^+^cells, stained for CD 172a, CD 14 and CD 16 and stimulated with Pam2CSK4, Poly(I:C), LPS, Gardiquimod, or Resiquimod for 15 min, before being fixed/permeabilized and stained with a fluorochrome-conjugated monoclonal antibody against phosphorylated p38 MAPK. Incubation with PBS served as control. Graphs show the fold change in median fluorescence intensity (MFI of stimulated sample divided by MFI of PBS control) of phospho-p38 MAPK staining for cM (CD14^high^CD16^−^), intM (CD14^high^CD16^high^) and ncM (CD14−CD16^high^). Data of four different animals (color-coded dots) is shown. Boxes indicate minimum, maximum and median values.

Bovine cM were clearly enriched in transcripts involved in antibacterial responses (Figure3B). These transcripts code for the accessory protein for TLR4 (*MD2*), other LPS-binding proteins (*AOAH (47)*, *CRISPLD2*(48), beta-defensins (*DEFB1*, *DEFB3*, *DEFB6*, *DEFB7*, *DEFB10*, *DEFB113*) (49), and other proteins commonly known or described to be involved in antibacterial responses (*RNASE6* (50), *BPI* (51, 52), *HP* (53), *CHI3L1* (54), *LYZ*). Notably, high levels of *MD2* were also expressed by intM of two animals. Three genes associated with antibacterial responses were found to be upregulated in ncM and intM – *ACOD1*, *CD180* and *LY86* (MD1), the latter two genes coding for LPS-binding proteins and members of the TLR family that form a complex to regulate TLR4 signaling (55). ACOD1 (IRG1) mediates the production of itaconate, which is – apart from its anti-inflammatory functions –also known for its antibacterial properties (56).

Looking at genes associated with antiviral responses, we found one gene upregulated in cM (*PTPN22*) and the vast majority of genes upregulated in intM and ncM (Figure 3C). PTPN22, overexpressed in cM, has been reported to potentiate TLR-induced type-I interferon production (57), and to regulate inflammasome activation (58). Both ncM and intM clearly expressed the highest levels of *IRF4* and *IFNAR2*, whereas *IFNAR*1 was expressed to similar levels in all monocyte subsets (data not shown). Also *IFNGR1* and *IFNGR2*, coding for the IFN-γ receptor, showed increased transcription in ncM and intM. Accordingly, the transcription of interferon-induced antiviral genes (*ISG15*, *RSAD2*, *IFIT1*, *IFIT2*, *IFI47*) was higher in ncM and intM, as compared to cM. Notably, the ubiquitin-like protein ISG15 exerts its antiviral function intracellularly by ISGylation of viral proteins and also extracellularly by acting in a cytokine-like manner to promote IFN-γ production of NK cells and T cells (59). Viperin (*RSAD2*), a multifunctional antiviral factor (60) has recently gained attention, as it was shown to act as a synthase for antiviral ribonucleotides (61).

Moreover, ncM and intM were enriched in transcripts for several interferon-induced guanylate-binding proteins (GBP1, GBP4, GBP5, GBP6, LOC107131333). Alongside the GBP4 gene displayed in the heatmap (ENSBTAG00000037634), also three other genes annotated as GBP4 (ENSBTAG00000014529, ENSBTAG00000038233) or GBP4-like (ENSBTAG00000002416) were upregulated in ncM and intM (data not shown). Only recently, GBP1 has been allocated an important role in apoptosis and pyroptosis of human macrophages (62). GBP4 has been reported to negatively regulate virus-induced type I IFN responses by targeting interferon regulatory factor 7 (63). Along this line, BST2, exclusively expressed in ncM and intM, was reported to inhibit type I interferon and cytokine production in TLR7/9-stimulated pDC (64). Furthermore, SEC14L1, more than 2-fold enriched in ncM and intM, was reported to negatively regulate RIG-I-mediated signaling (65).

Responsiveness of monocyte subsets to TLR stimulation, as determined by phosphoflow cytometry for p38 MAPK (Figure 3D), corroborates a specialization of cM and ncM for antibacterial and antiviral responses, respectively. While cM and intM showed the highest activation by the TLR4-ligand LPS, stimulation with the TLR7/8 ligands Gardiquimod and Resiquimod led to the strongest response in ncM. Also, responses to Pam2CSK4, a synthetic diacylated lipopeptide and ligand for TLR2/6, were highest in ncM. At last, similar low-level responses to Poly(I:C) were detectable in all subsets, with one animal standing out by a considerably higher response. Taken together, these results indicate that bovine monocyte subsets are specialized in responding to different pathogen-associated molecular patterns, suggesting a role of ncM in immunity against viruses (TLR7/8) as well as gram-positive bacteria and mycoplasma (TLR2/6) (66) and for cM and intM against gram-negative bacteria (TLR4).

### 3.4 Nonclassical monocytes have a gene expression signature promoting resolution of inflammation and tissue repair

All three monocyte subsets expressed anti-inflammatory genes, but ncM and intM were clearly dominant in this regard (Figure 4A).

**Figure 4.**
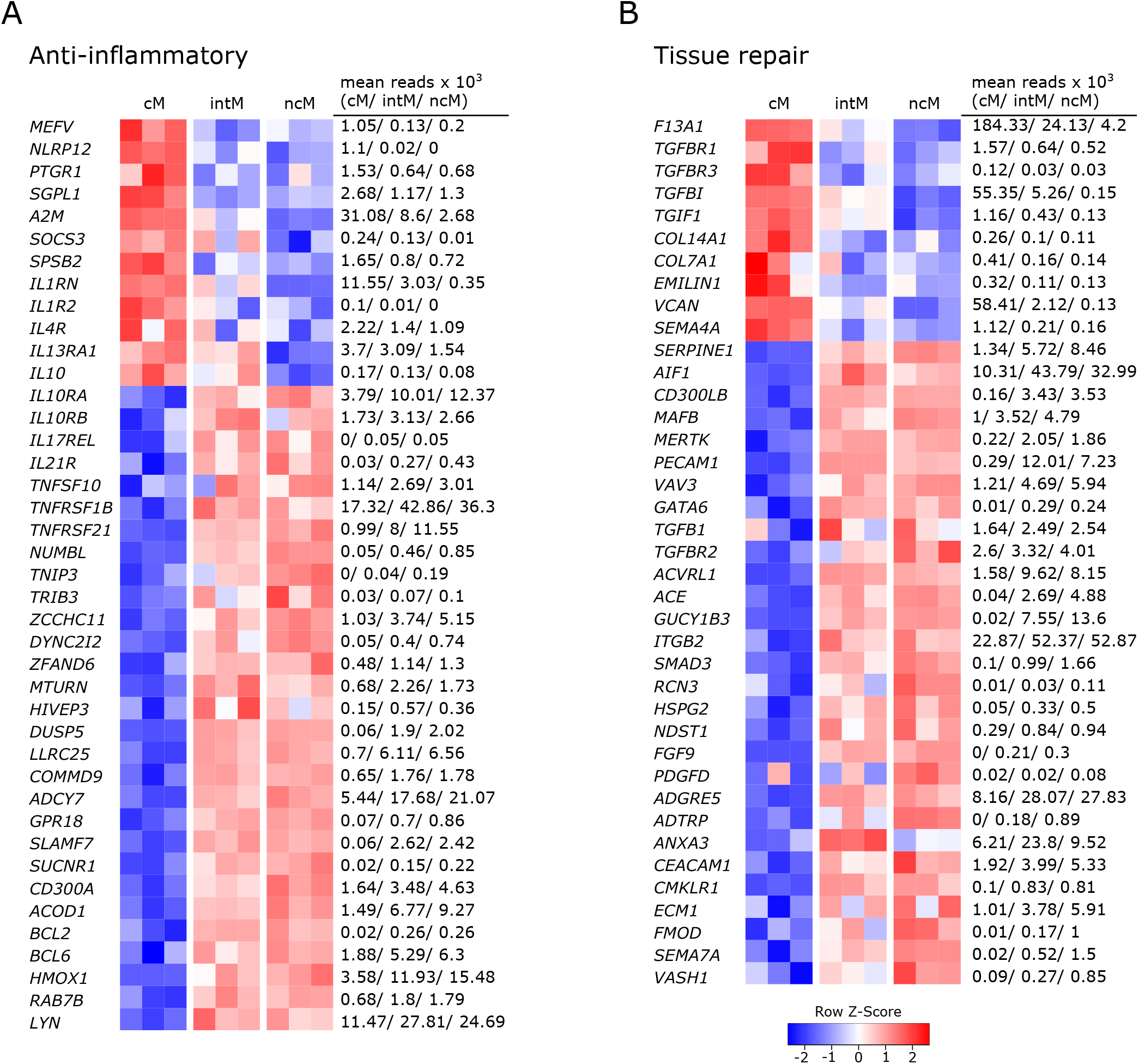
Gene expression associated with anti-inflammatory responses and tissue repair. Illumina sequencing was performed on RNA isolated from sorted monocyte subsets (cM, intM, ncM) of three animals. Heatmaps show row z-scores calculated from log2-transformed normalized counts of selected genes associated with anti-inflammatory responses (A), and tissue repair (B). Mean kilo reads for each subset and gene are given to the right of each heatmap.

With the expression of anti-inflammatory genes, cM seemed to mainly regulate their own pro-inflammatory functions. Among those genes are regulators of inflammasome activation (*MEFV* (67), *NLRP12* (68)), an enzymatic inactivator of leukotriene B4 (*PTGR1*) and sphingosine-1-phosphate (*SGPL1*) (69), a cytokine-scavenging protein (*A2M*) (70), as well as a negative regulator of cytokine signaling (*SOCS3*) and nitric oxide production (*SPSB2*). Moreover, cM were strongly enriched in transcripts for the IL-1 receptor antagonist (*IL1RN*) and exclusively expressed the decoy receptor for IL-1 (IL1R2). Also *IL4R* and *IL13A1* were expressed to higher levels in cM, suggesting that cM are especially receptive for IL-13, which was shown to inhibit the production of pro-inflammatory cytokines in macrophages (71). Furthermore classical signaling through the IL-6 receptor (*IL6R*), 3-fold enriched in cM over ncM (Figure 2A), is reported to mediate anti-inflammatory effects of IL-6, as opposed to signaling through soluble IL-6 receptor (28). Notably, while cM expressed the highest levels of *IL10*, transcripts for the IL-10 receptor (*IL10RA*, *IL10RB*) were clearly enriched in ncM and intM. Additionally, both ncM and intM contained *IL17REL* transcripts, coding for a soluble receptor and potential negative regulator of IL-17 signaling (72). Moreover, ncM and intM were enriched for *IL21R* transcripts, with IL-21 signaling described to enhance SOCS gene expression and to limit cytokine production in human monocyte-derived cells (73). Also transcripts for TRAIL (*TNFSF10*), TNFR2 (*TNFRSF1B*) and *TNFRSF21* were strongly overexpressed in ncM and intM. Apoptosis-inducing TRAIL expression on monocytes is suggested to be critical for regulation of inflammation (74), and clear overexpression of TNF receptor 2 (*TNFRSF1B*) in intM and ncM may favor suppressive signaling of transmembrane TNF-α, as reported for murine myeloid-derived suppressor cells (75). Death receptor 6 (*TNFRSF21*) has been reported to have inhibitory effects on monocyte differentiation when cleaved from the surface of tumor cells by matrix metalloproteinase 14 (*MMP14*) (76), the latter being about 10-fold overexpressed in ncM and intM (data not shown).

Other anti-inflammatory genes over-expressed by ncM and intM are reported to be mostly involved in negative regulation of NF-κB signaling (*NUMBL*, *TNIP3*, *TRIB3* (77), *ZCCHC11*, *DYNC2I2* (78), *ZFAND6* (79), *MTURN* (80), *HIVEP3* (81), *DUSP5*, *LRRC25*, *COMMD9* (82), *ADCY7* (83)), but also include surface receptors involved in regulation of inflammation, such the receptor of resolvin D2 (*GPR18*) (84), *SLAMF7* (85), *SUCNR*1 (86), and *CD300A* (87).

Notably, expression of *ACOD*1 was over 6-fold higher in ncM compared to cM. The metabolite itaconate, generated by IRG1 (ACOD1), is well-known for its anti-inflammatory effects (56, 88). Furthermore, *BCL2* and *BCL6* were transcribed to higher levels in ncM and intM. BCL-2 was shown to negatively regulate caspase-1 activation (89) and BCL-6 was recently reported to exert anti-inflammatory effects by suppressing IL6 transcription in murine macrophages (90). Like in human CD16^+^ monocytes (91), *HMOX1* was significantly increased in ncM and intM. Heme oxygenase-1 was shown to be induced by IL-10 and to mediate the anti-inflammatory effect of IL-10 in murine macrophages, presumably via NF-κB suppression by the heme degradation product carbon monoxide (92). Also in human monocytes, HMOX1 was reported to inhibit LPS-induced TNF-α and IL1-β production (93). Additionally, *RAB7B*, described to promote degradation of TLR4 (94) and TLR9 (95), was almost 2-fold higher expressed in ncM and intM. As was *LYN*, also expressed in human ncM and intM (96, 97), and reported to negatively regulate TLR-induced cytokine responses (98). Notably, LYN has recently been proposed as a negative regulator of murine ncM development (99).

Overall, these results indicate anti-inflammatory functions for ncM and intM.

In line with their suggested anti-inflammatory and pro-resolving functions, a number of genes associated with different stages of tissue repair were upregulated in ncM and intM (Figure 4B). While cM were enriched in *F13A1*, mediating hemostasis (100), intM and ncM expressed *SERPINE1*, described to regulate clot resolution (101). Both ncM and intM contained the highest transcript levels of genes associated with efferocytosis (*AIF1* (102), *CD300LB (103)*, *MAFB (104)*, *MERTK* (105), *PECAM1(106)*, *VAV3* (107)). Notably, MERTK has also been described as a negative regulator of human T-cell activation (108).

Furthermore, ncM expressed higher levels of *TGFB1* and of genes coding for TGF receptors (*TGFBR2*, *ACVRL1*) or being directly or indirectly involved in TGF-β pathways (*ACE*, *GUCY1B3*, *ITGB2*, *SMAD3*). Transforming growth factor beta (TGF-β) is known as a pro-fibrotic cytokine involved in wound healing (109). Notably, cM expressed higher levels of the TGF receptor genes *TGFBR1* and *TGFBR3*, and of *TGFBI* and *TGIF1*. Genes associated with extracellular matrix components dominantly transcribed by ncM included *HSPG2* (basement membrane-specific heparin sulfate) and *NDST1* (biosynthesis of heparin sulfate), whereas cM contained the highest transcript levels for collagens *COL14A1*, *COL7A1*, as well as *EMILIN1* and *VCAN*. Furthermore, fibroblast growth factor *FGF9* and platelet-derived growth factor *PDGFD* were clearly enriched in ncM. The high expression of *CMLKR1* in intM and ncM suggests that these subsets are attracted to inflamed tissues via chemerin and contribute to resolution of inflammation via binding to resolvin E1 (110), which was shown to increase IL-10 production and phagocytosis of apoptotic neutrophils in macrophages (111, 112).

In addition, a number of metalloproteinases and their regulators were either upregulated in ncM (*ADAM9*, *ADAMTSL5*, *ADAMTS12*, *MMP14*, *MMP19*, *TIMP2*, *TSPAN14*, *ECM1*) or cM (*ADAM19*, *ADAM8*, *ADAMTS2*, *MMP25*) (data not shown). Several metalloproteinases are described to be involved in wound healing (113), among which MMP14 and ADAM9 are suggested to regulate epithelial cell proliferation (113, 114). Finally, several genes associated with angiogenesis (*ADGRE5*/CD97 (115), *ADTRP*, *ANXA3 (116)*, *CEACAM1 (117)*, *CMKLR1* (118), *ECM1 (119), FMOD (120)*, *SEMA7A (121)*, *VASH1*) were predominantly expressed by intM and ncM. Notably, *SEMA4A* (122), also reported to be involved in angiogenesis, showed the highest transcription in cM. Altogether, these data suggest anti-inflammatory pro-resolving functions of ncM and intM, as well as a prominent role of these subsets in wound healing and tissue regeneration.

### 3.5 Gene expression indicates differential capabilities for antigen presentation, co-stimulation, and modulation of T-cell responses

Antigen presentation capabilities are reported for monocytes across species (1). As expected from phenotypic analyses, intM stood out by their high gene expression for MHC-II (*BOLA-DQA5*, *BOLA-DQB*, *BOLA-DRA*), and the co-stimulatory molecules CD40 and CD86 (Figure 5A). Interaction of CD40 with CD40L on T cells has been reported to stimulate Th17 responses (123). Notably, genes for MHC class I molecules (BOLA, BoLA), were transcribed to higher levels in intM and ncM. Furthermore, mRNA from genes associated with the presentation of lipid antigens to T cells (*LOC515418* annotated as CD1a molecule-like, *CD1E*), was enriched in intM.

**Figure 5.**
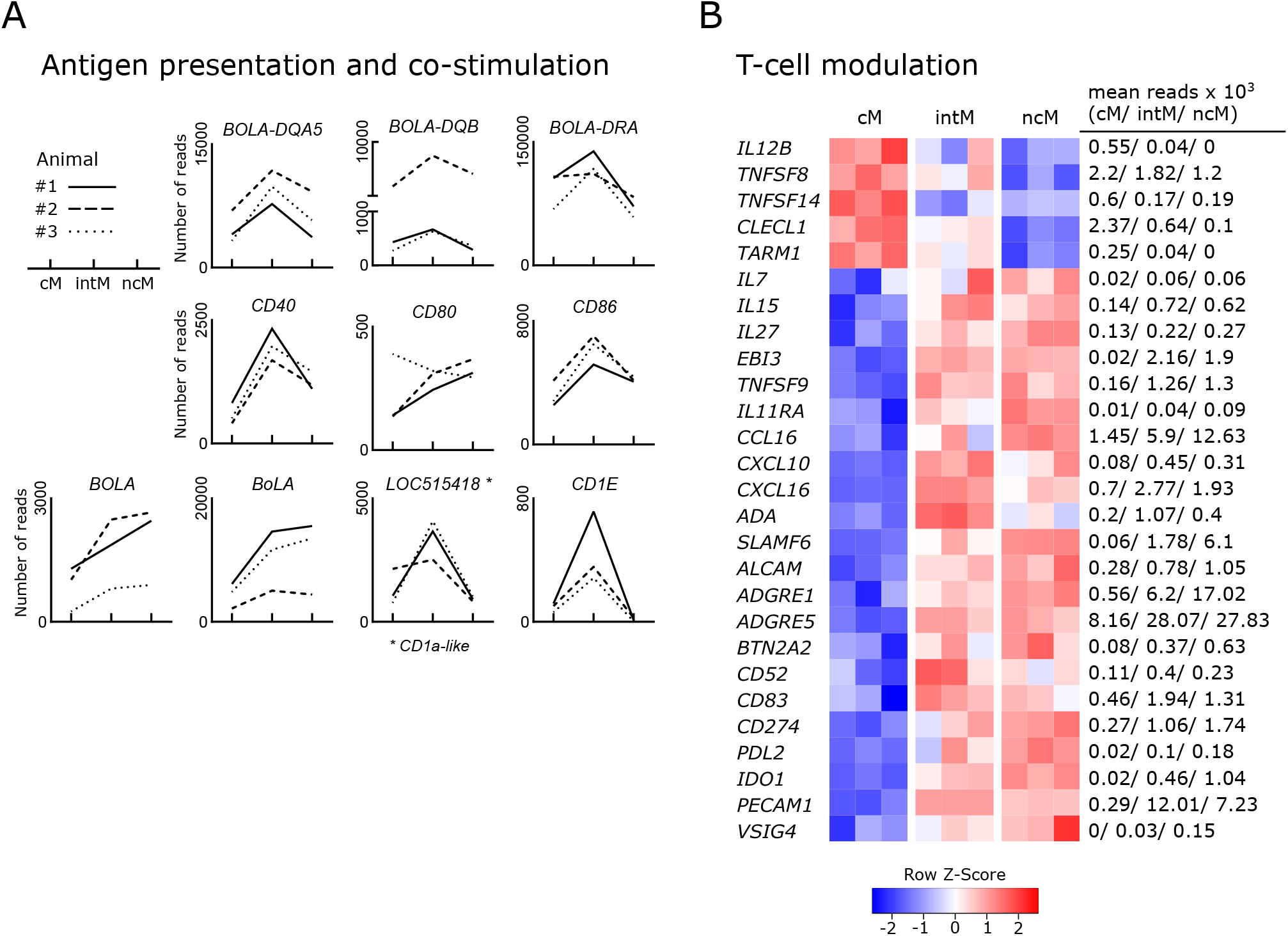
Expression of genes involved in the shaping of T-cell responses. Illumina sequencing was performed on RNA isolated from sorted monocyte subsets (cM, intM, ncM) of three animals (#1-3). A) Gene expression promoting antigen presentation and co-stimulation. Graphs show the number of reads across monocyte subsets for selected genes with individual animals indicated by solid (#1), dashed (#2) and dotted (#3) lines. (B) Gene expression involved in T-cell modulation. Heatmap shows row z-scores calculated from log2-transformed normalized counts of selected genes. Mean kilo reads for each subset and gene are given to the right of the heatmap.

Genes encoding T-cell signaling cytokines predominantly expressed by cM included *IL12B* (Figure5B), as well as *TNFSF8* (CD30L) and *TNFSF14* (LIGHT) (124, 125). In fact, transcription of *IL12B* mRNA was found to be absent in ncM, and over 13-fold increased in cM over intM. Two further genes upregulated in cM and reported to be involved in regulation of T-cell activation include *CLECL1* and *TARM1*. *CLECL1*, 20-fold increased over ncM, has been reported to act as a T-cell costimulatory molecule, skewing the CD4 T-cell response towards Th2 by increasing IL-4 production and proliferation (126). *TARM1*, about 4-fold enriched in cM, has been reported to suppress CD4-T-cell activation and proliferation *in vitro* (127).

T-cell signaling cytokines predominantly expressed by intM and ncM included *IL7* and *IL15*, reported to function in lymphoid homeostasis (128), and *IL27*, encoding a multifaceted cytokine described to both promote and suppress T-cell responses (129). Also, *EBI3* (IL-27β), an essential component of the cytokines IL-27 and IL-35, was exclusively expressed in intM and ncM (approx. 110-fold increased). Moreover, *TNFSF9*, a ligand for CD137 on T cells shown to be important for the generation of antiviral CD8-T-cell responses (130), was 8-fold enriched in ncM over cM.

Furthermore, the receptor for IL-11 (*IL11RA*) was expressed at higher levels in ncM and intM. In line with almost absent *IL12B* transcription in ncM and intM, IL-11 signaling has been reported to inhibit IL-12 production in macrophages (131), which supports polarization towards Th2 responses (132). As shown in Figure 2A and Figure 5B, chemokines relevant for T-cell responses were mainly overexpressed by ncM and intM (*CCL3*, *CCL4*, *CCL5* and *CCL16*, *CXCL10*, *CXCL16*).

In fact, the vast majority of genes associated with modulation of T-cell responses was overexpressed in ncM and intM. Intermediate monocytes expressed the highest levels of adenosine deaminase (*ADA*), which is reported to act as a modulator of T-cell differentiation, increasing the generation of effector, memory and regulatory T cells (133). Both ncM and intM were enriched in transcripts for *SLAMF6*, reported to boost IFN-γ production and cytolytic anti-tumor activity of human CD8 T cells *in vitro* (134). Furthermore, expression of *ALCAM*, encoding a ligand for CD6 on T cells, important for stabilizing the immunological synapse between APC and T cells (135) and reported to mediate extravasation of monocytes (136), was more than 3-fold higher expressed in ncM than in cM. Notably *ADGRE1* (F4/80), reported to be essential for the generation of Re and peripheral tolerance when expressed on antigen-presenting cells (137), was 28-fold enriched in ncM over cM. Moreover, *ADGRE5*, coding for CD97 and described to induce regulatory T cells and IL-10 production upon engagement of CD55 on CD5 T cells (138, 139) was 3-fold higher transcribed in ncM and intM. Consistent with the idea that ncM and intM promote the generation of Tregs, they showed an almost 8-fold higher transcription of *BTN2A2*, a butyrophilin reported to inhibit activation and induce Foxp3 expression in murine T cells (140, 141).

Many T-cell modulating genes overexpressed in ncM and intM were found to be genes involved in negative regulation of T-cell activation (*BTN2A2*, *CD52*, *CD83*, *CD274*/PDL1, *PDL*2, *IDO1*, *PECAM1*, and *VSIG4*). Soluble CD52 was reported to suppress T-cell activation via binding to Siglec-10 (142) and soluble CD83 was shown to inhibit T-cell activation by binding to the TLR4/MD-2 complex on human CD14^+^ monocytes and inducing expression of anti-inflammatory mediators such as IDO and IL-10 (143). Transcription for PDL-1, a well-known inhibitor of T-cell activation (144), was increased 5-fold in ncM over cM. Notably, PDL-1 has recently been employed as a marker of ncM for *in-vivo* tracking of this monocyte subset in mice (145). Similarly, the gene for PDL-2, a second ligand for PD-1 on T cells with T-cell inhibitory function (146), was 9-fold higher expressed in ncM, though at lower levels than the gene for *CD274* (PDL-1). Strikingly, *IDO1* expression was found to be significantly increased in intM (20-fold) and ncM (30-fold) when compared to cM, where expression was almost absent (mean of 20 reads). IDO1 was described to inhibit T-cell activation by degrading tryptophan, and to promote tolerance of DC and the expansion of Tregs (147). Also transcription of *PECAM1* (CD31), described as a key co-inhibitory receptor promoting tolerogenic functions in both DCs and T cells through homophilic interactions (148), was greatly upregulated in ncM and intM. A recent *in vitro* study also suggested that high CD31 expression on DCs reduces priming of CD4 T cells by impairing stable cell-cell contacts (149). Furthermore, *VSIG4*, coding for a B7 family-related protein specifically expressed on resting macrophages was primarily expressed in ncM. Notably, VSIG4 has been described as a strong negative regulator of T-cell activation, maintaining T-cell unresponsiveness in healthy tissues (150),

Taken together, these data clearly indicate that monocyte subsets are actively involved in the shaping of T-cell responses, with ncM and intM being especially well equipped for T-cell suppression, either directly or via the induction of regulatory T cells.

### 3.6 Expression of metabolic genes differs markedly between classical and nonclassical monocytes

Given that metabolism and immune function are tightly linked (151–153), differences in metabolic pathways can give indications for subset-specific functions. In fact, monocyte subsets showed prominent differences in the expression of genes relating to metabolism. The vast majority of these differentially expressed genes was associated with glycolysis and was overexpressed in cM. The expression of glycolysis-related genes is illustrated in Figure 6A. The transcription factor *HIF1A* is shown on top, followed by glycolytic enzymes ordered from beginning to end of the glycolytic pathway. Apart from glycolytic genes, also genes involved in oxidative phosphorylation showed increased transcription in cM as compared to ncM (Figure 6B). Among those genes was the subunit of the mitochondrial ATP synthetase (*ATP5I*), a mitochondrial inner membrane protein (*MPV17*) supporting oxidative phosphorylation (154), the genes coding for the components of complex II of the mitochondrial electron transport chain succinate dehydrogenase (*SDHA*, *SDHB*, *SDHC*, *SDHD*), and also most of the numerous genes coding for subunits of complex I (data not shown). In line with increased glycolysis in cM, transcripts for the glucose transporters *SLC2A1* and *SLC2A3* were about 3-fold enriched in cM (data not shown). As reported in our previous publication (9), also SLC genes involved in transport of succinate (*SLC13A3*), citrate (*SLC13A5*) and lactate/pyruvate (*SLC16A1*) were mainly expressed in cM. The metabolites succinate and citrate are both described as critical pro-inflammatory mediators linking metabolism to immune functions (155). Moreover, two genes associated with fructose metabolism (*KHK*, *SORD*) were overexpressed in cM. Fructose-induced metabolic changes were recently described to enhance inflammatory responses of dendritic cells (156). In addition, genes involved in the oxidative (*G6PD*, *PGD*) and non-oxidative (*TALDO1*) pentose-phosphate pathway (PPP) showed the highest transcription in cM. The PPP, providing redox-equivalents and nucleotide precursors, was shown to be essential for pro-inflammatory functions of human macrophages (157, 158). Furthermore, *PANK1*, coding for a key enzyme in CoA synthesis was 5-fold upregulated in cM.

**Figure 6.**
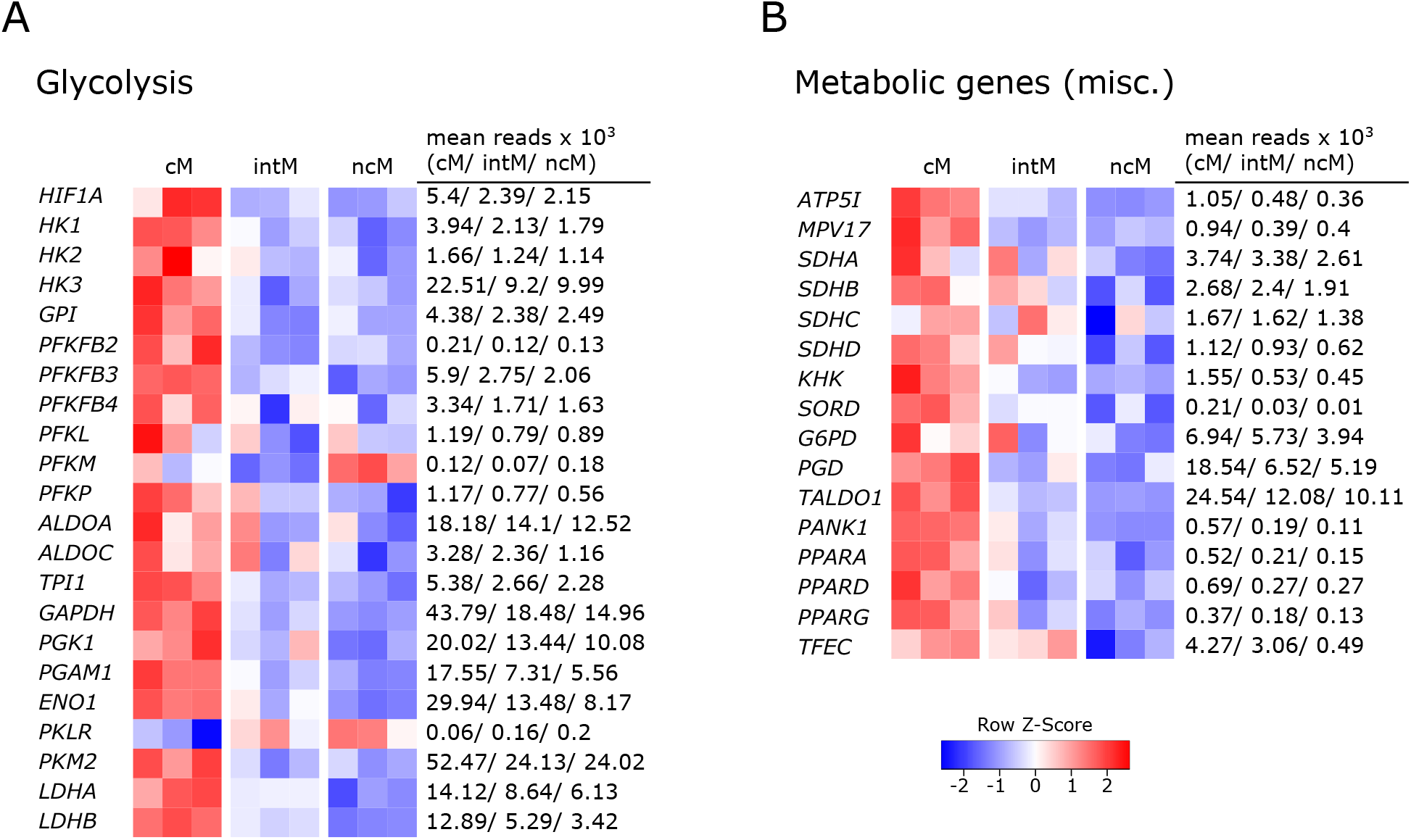
Metabolic gene expression. Illumina sequencing was performed on RNA isolated from sorted monocyte subsets (cM, intM, ncM) of three animals. Heatmaps show row z-scores calculated from log2-transformed normalized counts of glycolytic genes (A) and of genes associated with other metabolic pathways (B). Mean kilo reads for each subset and gene are given to the right of each heatmap.

Notably, all three genes coding for members of the PPAR family (*PPARA*, *PPARD*, *PPARG*) were transcribed at higher levels in cM. These lipid-activated nuclear receptors have evolved as key regulators linking lipid metabolism to inflammation, and in particular expression of PPARG is associated with anti-inflammatory functions (159). We recently showed that bovine cM downregulate *PPARG* transcription dramatically upon Gardiquimod (TLR7/8) stimulation *in vitro* (16). PPARA was recently proposed as an important mediator of antimicrobial responses to mycobacteria (160), inducing expression and translocation of TFEB, a key transcriptional activator of autophagy and lysosomal biogenesis. Notably, while *TFEB* was equally expressed in all monocyte subsets (data not shown), transcription of *TFEC*, a less well described member of the TFE family, was 10-fold increased in cM and intM compared to ncM (mean reads: 4000/3000 vs. 400).

Altogether, these data suggest that cM are metabolically more active than ncM, with a significantly enhanced expression of genes involved in glycolysis, supporting pro-inflammatory functions.

### 3.7 Mitochondrial respiration prevails in nonclassical monocytes

Having observed an upregulation of a variety of metabolic genes in cM, we were interested in the actual contribution of glycolysis and mitochondrial respiration to ATP production in the different monocyte subsets (Figure 7A). Extracellular flux analysis of sorted monocyte subsets demonstrated that ncM predominantly used oxidative phosphorylation (OXPHOS) for ATP production, whereas the proportion of ATP produced by glycolysis was higher in intM and highest in cM (Figure 7B+C).

**Figure 7.**
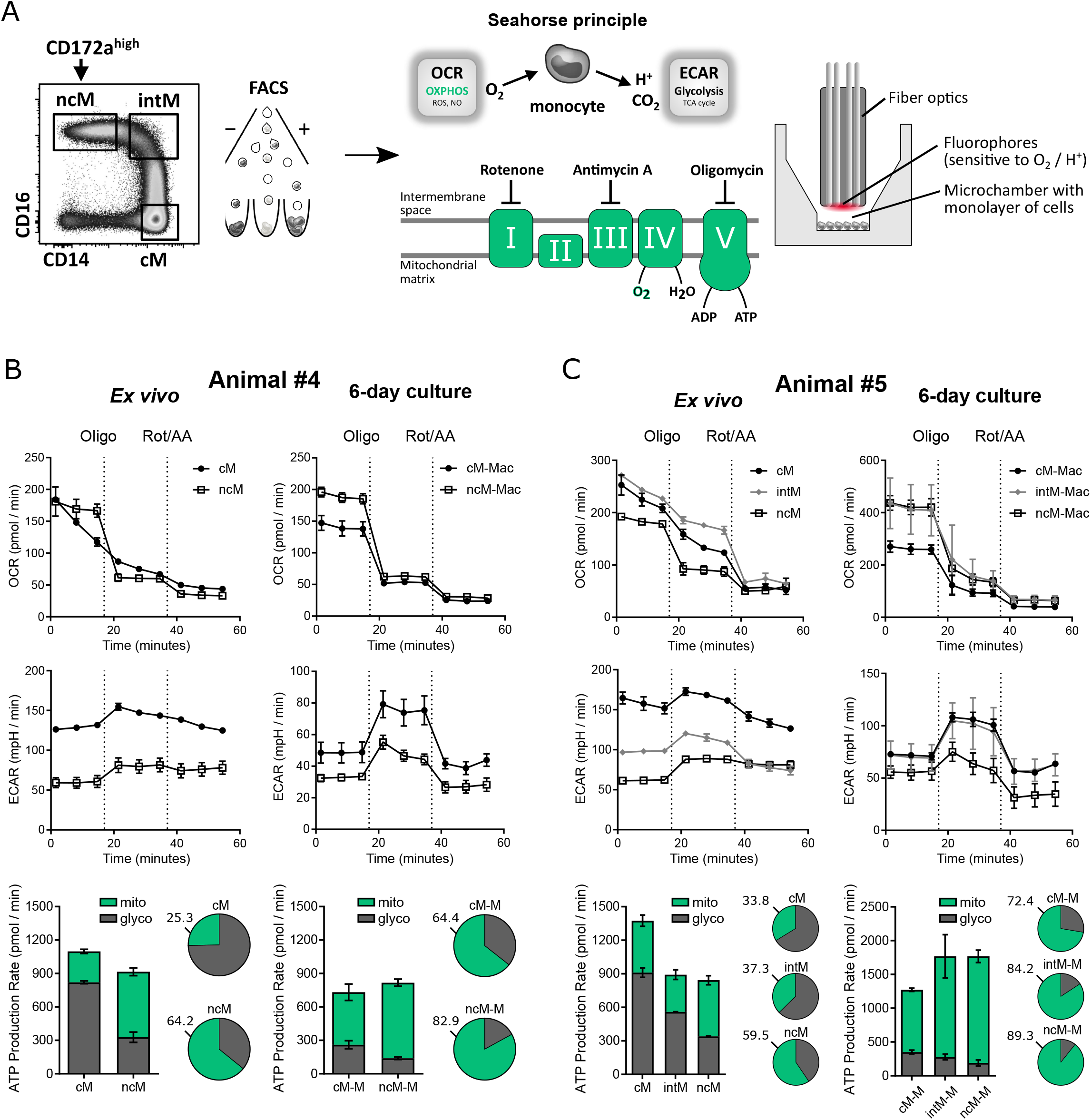
Mitochondrial respiration prevails in nonclassical monocytes. Bovine monocyte subsets were FACS-sorted and metabolic activity was analyzed by Agilent Seahorse XF technology and the XF Real-Time ATP Rate Assay. Oxygen consumption rate (OCR) and extracellular acidification rate (ECAR) were automatically calculated from measured oxygen and pH decrease (Agilent Wave software). The electron transport chain inhibitors oligomycin and rotenone/antimycin A were injected sequentially to allow calculation of OXPHOS-as well as glycolysis-mediated ATP production rates (mito, glyco) from resulting OCR and ECAR. Monocyte subsets from four animals were analyzed (two animals with intM, and three animals after 6-day *in vitro* culture). Representative data from two animals are shown. (A) Gating strategy as well as principle of Agilent Seahorse XF technology is illustrated. (B+C) OCR and ECAR traces as well as ATP production rates as absolute values (bar graphs) and relative values (pie charts) are shown for cM and ncM seeded in triplicates (B), or for all three monocyte subsets (cM, intM, ncM) seeded in duplicates (C). For cM of animal #4, only two out of three wells were included in the analysis. Macrophages derived from monocyte subsets after six days of culture in the presence of M-CSF are shown in the right panels of B and C.

After 6 days of *in-vitro* culture, the proportion of ATP produced by OXPHOS increased in all subsets. Generally, the metabolic activity was considerably higher in monocyte-derived macrophages (cM-Mac, intM-Mac, ncM-Mac), which is why the optimal cell number for seahorse assays had to be reduced by five times. The proportional increase in OXPHOS-generated ATP was most evident in intM-Mac and cM-Mac. However, ncM-Mac remained to be the subset with the highest contribution of OXPHOS. Overall, the observed preferences for different metabolic pathways further support the diverging roles of monocyte subsets in inflammation and beyond.

## 4 Discussion

With the present study, we extended the phenotypic characterization of bovine monocyte subsets, and combined an in-depth analysis of their transcriptomes with metabolic and TLR-stimulation assays in order to get a detailed insight into subset-specific functions. Pairwise comparison of gene expression coupled with extensive literature research has revealed substantial transcriptomic differences between bovine monocyte subsets, likely determining their specializations.

Bovine cM clearly emerged as pro-inflammatory, with overall gene expression supporting antibacterial inflammatory responses. This is in line with data on human and murine cM, and with earlier studies of bovine cM that have shown their superiority in phagocytosing bacteria (27). Both ncM and intM were dominant in the transcription of many genes associated with regulatory functions. The expression of anti-inflammatory genes and numerous genes associated with wound healing (efferocytosis, angiogenesis, fibrosis) clearly indicate that bovine ncM are specialized in the resolution of inflammation and tissue regeneration, as suggested for ncM based on studies in mouse models (22). In line with our steady-state transcriptomic data, bovine ncM were shown to almost lack IL-1β production upon inflammasome activation *in vitro* (27). However, literature on the ability of human ncM to produce IL1-β is conflicting (6, 161). As reported for human intM (6), the majority of genes in bovine intM was found to be expressed at levels in-between cM and ncM, while showing the highest transcriptional similarity with ncM. Notably, bovine intM were reported to produce the highest amounts of reactive oxygen species in response to opsonized bacteria and the highest amounts of IL-1β following inflammasome activation (27). This is surprising when looking at the steady-state transcriptome of intM described in the present study.

Literature on intM is conflicting (18), one reason being that human intM and ncM were often analyzed as one CD16^+^ subset in clinical studies. Another explanation for observed discrepancies may be that cM upregulate CD16 upon activation, which makes them phenotypically indistinguishable from intM when using the standard gating strategy with CD14 and CD16. As reported recently, an upregulation of CD16 on human cM was mistaken for an expansion of intM in response to dengue-virus infection (162). Also for bovine cM, an upregulation of CD16 is reported following stimulation with IFN-γ (27). Moreover, we found that sorted bovine cM (CD14^+^CD16^−^) were all CD16^+^ after overnight culture, even without stimulation (unpublished observation). It is therefore likely that *bona fide* intM represent a different subset than CD14^+^CD16^+^ cells reported to increase following infection. And functional assays may indicate pro-inflammatory capabilities of intM because of contamination with highly activated cM in the intM gate. Therefore, also dominant glycolysis and LPS responsiveness of intM, observed in the present study and both reminiscent of cM, should be interpreted in the light of possible gate contamination. An alternative gating strategy was recently proposed for human monocyte subsets (162), in order to avoid the problem of continuous CD14 expression and easily induced CD16 expression, also seen with bovine monocyte subsets. With the phenotyping at hand, it should be possible to establish an alternative gating strategy also for cattle. Transcriptomic analyses of these newly defined subsets should then provide clarity in regard to the poorly defined intM subset. In fact, heterogeneity of intM has been described for both humans (163) and mice (164). The observation that animal-to-animal variability was most prominent for intM in the current dataset, supports the idea that also bovine CD14^+^CD16^+^ intM are a mixed population.

Transcriptomic data also suggest diverging functions of bovine monocyte subsets in the interaction with T cells. Notably, intM showed the highest expression of MHC-II, both on mRNA and protein level. High expression of MHC-II is also reported for human intM (5, 6), and may be linked to their superiority in stimulating human CD4-T-cell proliferation (5). Considering also the high expression of CD86 and the high transcription of a CD1a-like gene, bovine intM may be particularly well equipped for co-stimulation and antigen presentation to T cells. Nonclassical monocytes and intM were enriched in transcripts for various genes promoting CD8-T-cell responses. This preferential activation of CD8 T cells and the danger associated with uncontrolled cytolytic T-cell responses might also explain why ncM and intM highly express genes mediating the inhibition of T cells and the generation of Tregs, the latter being reported for murine ncM (165). Furthermore, gene expression promoting activation of CD8 T cells, together with the interferon-associated gene signature and the high responsiveness to Gardiquimod and Resiquimod, indicate a specialization of bovine ncM towards antiviral responses, as suggested for human ncM (21). In contrast to ncM and intM, which almost lacked CD62L expression on protein level, high CD62L expression on bovine cM should allow this subset to enter lymph nodes directly from blood. Furthermore, lymph-mediated entry of antigen-presenting monocytes has been reported for mice (166). However, given the lack of CCR7 upregulation on stimulated bovine monocytes (16), the mechanisms that would enable this migration remain elusive.

Pro- and anti-inflammatory functions of cM and ncM, respectively, are also supported by their metabolic transcriptome, clearly indicating that cM are metabolically more active and skewed towards glycolysis. Differential use of ATP-generating metabolic pathways could also be confirmed by extracellular flux analysis, where cM predominantly performed glycolysis and ncM mainly employed oxidative phosphorylation (OXPHOS). After 6 days of culture and differentiation to macrophages *in vitro*, OXPHOS was strongly upregulated in cM-Mac and intM-Mac, but the fraction of ATP produced by OXPHOS was still highest in ncM-Mac. These observations are in line with the reported lower glucose uptake by bovine ncM (167). Apart from dominant OXPHOS, the anti-inflammatory nature of bovine ncM-Mac is supported by high transcription of *ARG1* (7, 168), encoding a molecule that is reported to suppress NK cells and T cells (169, 170). Overall, the metabolic preferences of bovine cM and ncM are in line with gene expression as well as respirometric measurements in human cells (171), suggesting similar metabolic programing and functional specialization of monocyte subsets in human and cattle.

## 5 Conclusions

Bovine monocyte subsets show clear subset-specific differences in regard to phenotype, function-related gene expression, metabolism, and TLR responsiveness. Transcriptomic analyses suggest pro-inflammatory antibacterial functions of cM, whereas ncM appear to be anti-inflammatory, antiviral, Treg- and CD8-T-cell supporting and specialized in tissue repair. A role of ncM in antiviral responses is further supported by their strong response to TLR7/8 stimulation. Although the transcriptome of intM showed high similarity with ncM, intM appeared closer to cM when looking at their metabolism and their TLR responsiveness. Metabolically, cM favor glycolysis, whereas ncM mainly perform oxidative phosphorylation. Differentiation to macrophages in the absence of stimulation massively induced OXPHOS in cM. Future studies are required to address how the metabolism of monocyte subsets is reprogrammed upon stimulation. Overall, the results of the present study are highly valuable for ensuing functional investigations of monocyte subsets across species.

## 6 Acknowledgments

We want to thank the team of the Clinic for Ruminants at the Vetsuisse Faculty in Bern for blood sampling, and Giuseppe Bertoni for obtaining the approval for it. We also want to thank Sylvie Python and Gael Auray for their support with cell sorting and RNA extraction and Corinne Hug for her help in preparing monoclonal antibodies. Furthermore, we thank Stefan Müller and Thomas Schaffer (FACSLab, University of Bern) for cell sorting. Finally, we thank Muriel Fragnière and Tosso Leeb from the Next Generation Sequencing Platform of the University of Bern for RNA sequencing, and Irene Keller and Rémy Bruggmann (Interfaculty Bioinformatics Unit, University of Bern) for performing initial bioinformatic analyses.

