## Supplementary Figure 1 for "Transcriptomic signature and metabolic programming of bovine classical and nonclassical monocytes indicate distinct functional specializations"

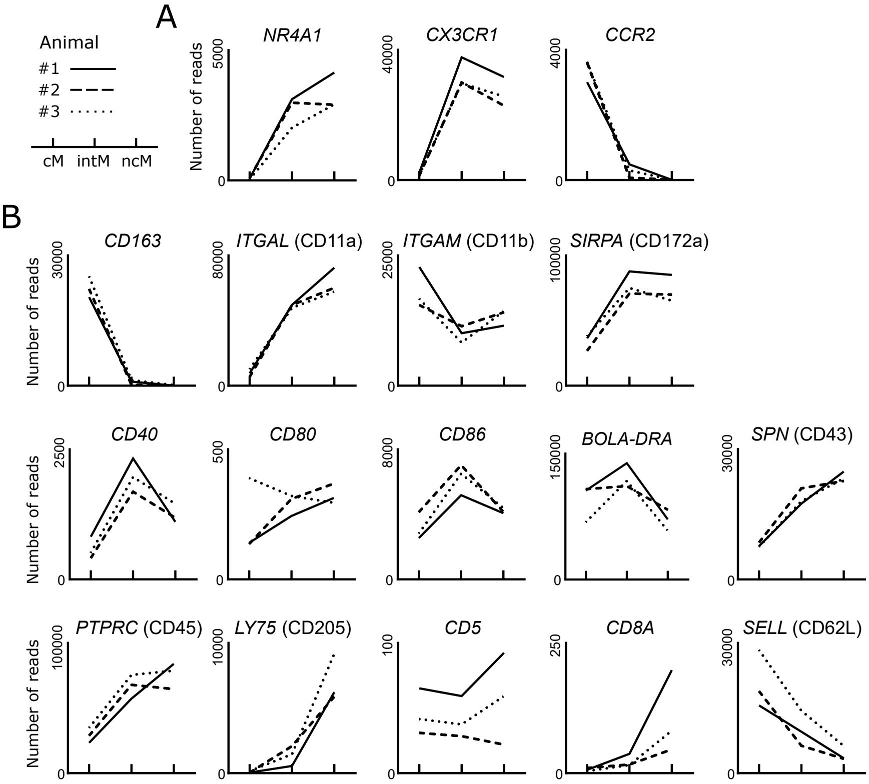


**Supplementary Figure 1** Transcription of key genes (A) and genes previously analyzed by flow cytometry (B). Illumina sequencing was performed on RNA isolated from sorted monocyte subsets (cM, intM, ncM) of three animals (#1-3). Graphs show the number of reads across monocyte subsets for selected genes with individual animals indicated by solid (#1), dashed (#2), and dotted (#3) lines.
